# The mutational signatures of poor treatment outcomes on the drug-susceptible *Mycobacterium tuberculosis* genome

**DOI:** 10.1101/2022.11.20.517260

**Authors:** Yiwang Chen, Qi Jiang, Mijiti Peierdun, Howard E. Takiff, Qian Gao

## Abstract

Drug resistance is a known risk factor for poor tuberculosis (TB) treatment outcomes, but the contribution of other bacterial factors to poor outcomes in drug susceptible TB is less understood. Here, we generate a population-based dataset of drug-susceptible *Mycobacterium tuberculosis* (MTB) isolates from China to identify factors associated with poor treatment outcomes. We sequenced the whole genome of 3496 MTB strains and linked genomes to patient epidemiological data. A genome-wide association study (GWAS) was performed to identify bacterial genomic variants associated with poor outcomes. Risk factors identified by logistic regression analysis were used in clinical models to predict treatment outcomes and their associations were assessed with structural equation models (SEM). GWAS identified fourteen MTB variants (24.2% vs 7.5%, P<0.001) and a *de novo* reactive oxygen species (ROS) mutational signature (26.3%±18.2% vs 22.9%±13.8%, P=0.027) that were more frequent in patients with poor treatment outcomes. Patient age, sex, and duration of diagnostic delay were also independently associated with poor outcomes. The best clinical prediction model, with an AUC of 0.74, incorporates both host and bacterial risk factors, and host factors are more important. Together, our results reveal that although host factors are the most important determinants for poor treatment outcomes, the genomic characteristics of the infecting MTB strain may also contribute significantly to poor treatment outcomes. Fourteen genetic variants were statistically associated with poor TB treatment outcomes, but the optimal model for predicting treatment outcomes includes both patient characteristics and bacterial genomic determinants.

## INTRODUCTION

Tuberculosis (TB), caused by *Mycobacterium tuberculosis* (MTB), is responsible for more deaths globally than any other single infectious agent except SARS-COV2. TB causes nearly 1.5 million deaths and an estimated 10 million new cases each year (World Health Organization, 2021). Successful treatment of TB not only cures the patient but prevents disease transmission and the development of difficult-to-treat drugresistant strains. Treatment outcomes are therefore important metrics for assessing the effectiveness of national TB control programs (Dheda et al., 2017; Migliori et al., 2019). Although approximately 86% of patients with drug-susceptible TB are cured with the standard four-drug treatment (World Health Organization, 2021), that leaves a substantial subset of patients who fail treatment. To formulate strategies to reduce treatment failures, it is necessary to first define the risk factors associated with poor outcomes.

There are host factors that are well known to be associated with treatment failure, including patient adherence (Alipanah et al., 2018), sex and age (Imperial et al., 2018), diagnostic delay (Lestari et al., 2020), co-infection with the human immunodeficiency virus (HIV) or TB treatment history (Bastos et al., 2017; Chenciner et al., 2021), but we wondered whether the genetic composition of the infecting strain might also contribute to poor outcomes. While drug resistance is the major bacterial risk factor for TB treatment failure (Lange et al., 2019; Mirzayev et al., 2021), a growing number of genomic studies suggest that there are other bacterial determinants that may also be risk factors. For example, specific mutations in metabolism-related genes of MTB could lead to drug tolerance (Hicks et al., 2018; Torrey et al., 2016), thus increasing the risk of developing resistance and relapsing after treatment (Brauner et al., 2016; Liu et al., 2020). Liu et al. (Liu et al., 2022) recently showed that genetic mutations in the transcriptional regulator *resR* gene cause antibiotic resilience that is associated with the acquisition of drug resistance and treatment failure. Although these genetic polymorphisms in the MTB genome do not directly lead to drug resistance, they are significantly more common in drug resistant bacteria (Hicks et al., 2018; Liu et al., 2022). Therefore, because drug resistance is a dominant risk factor for treatment failure, the association of other genomic determinants with treatment failure is best assessed in drug-susceptible TB isolates.

To study the role of bacterial genomic determinants other than canonical drug resistance that might be associated with poor treatment outcomes, we analyzed drug-susceptible TB isolates from new TB cases collected in population-based cohort studies at three different sites in China. The bacterial determinants identified and patient characteristics were then used to build a clinical prediction model to estimate the contribution of bacterial factors to poor TB treatment outcomes.

## RESULTS

### Characteristics of the study population and MTB isolates

The pooled population from the three different sites in China consisted of 3496 new cases with drug-susceptible TB. The patients were divided into three groups based on their treatment outcomes: good outcomes (88.8%, 3105/3496), poor outcomes (2.6%, 91/3496) and other (8.6%, 300/3496). To explore the bacterial factors associated with poor TB treatment outcomes, we first excluded patients with other outcomes, including lost to follow-up, non-TB deaths and unknown outcomes, because associations with bacterial factors would be unlikely. Ultimately, a total of 3196 new cases with drug-susceptible TB were included in this study (Figure 1A): 3105 with good outcomes and 91 with poor outcomes (failure, 25; TB death, 15; transferred for MDR, 4; and relapse, 47). The study patients were recruited from Shanghai (49.1%, 1569/3196), Sichuan (30.6%, 979/3196) and Heilongjiang (20.3%, 648/3196), China (Figure 1A). They had a mean age of 42.1 ± 18.2 years and 72.4% (2313/3196) were male. WGS was performed on all 3196 isolates, with an average depth of 100× and average genome coverage of 98%. Phylogenetic analysis of WGS data showed that nearly three quarters of the isolates were lineage 2 (74.2%, 2373/3196), with more than half belonging to the modern Beijing sublineage L2.3 (54.9%, 1754/3196) (Figure 1B).

**Figure 1.**
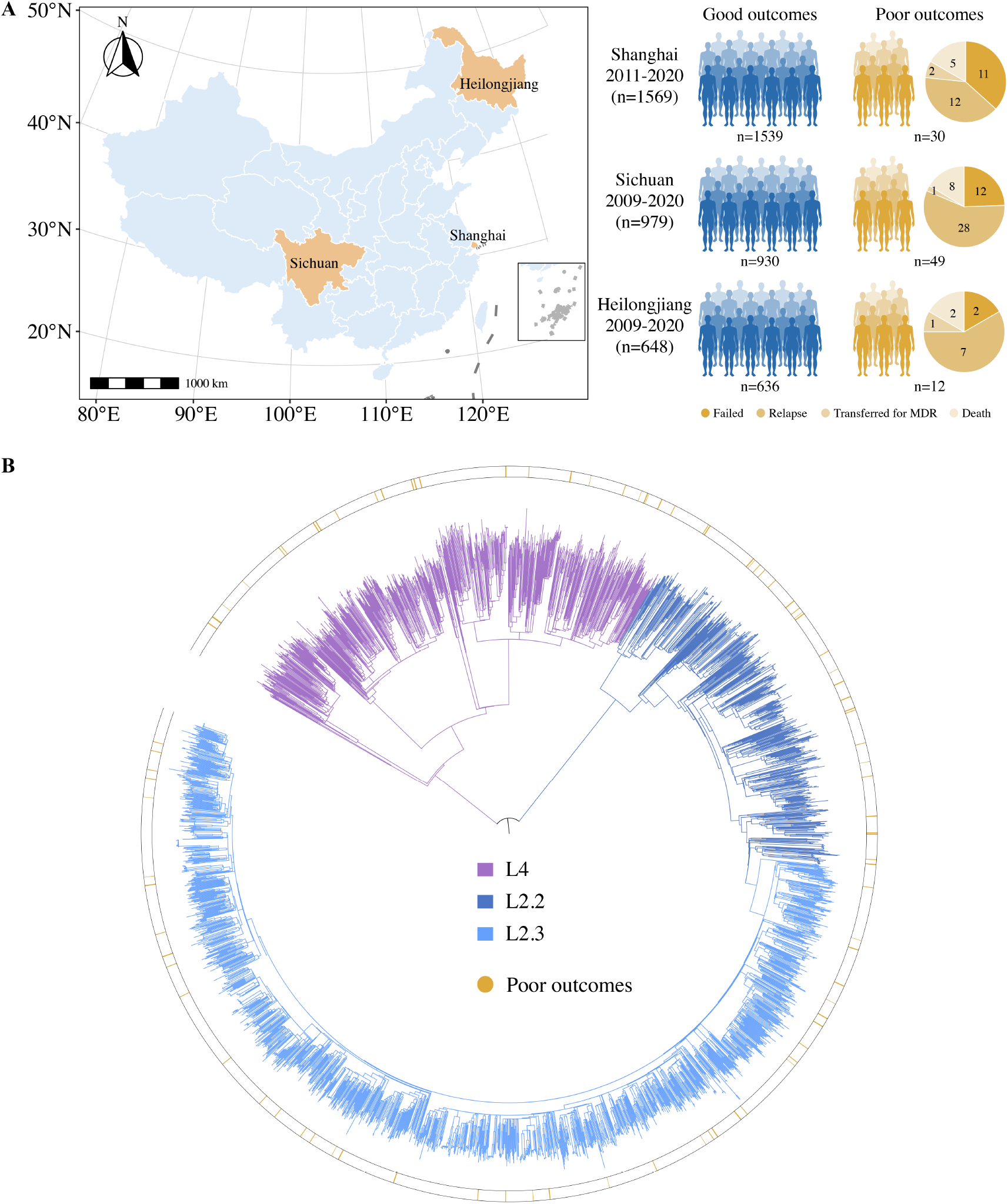
Sample origin and genetic structure of *Mycobacterium tuberculosis*. **(A)** Geographic location of the samples analyzed and study cohort characteristics. **(B)** The phylogenetic tree of 3196 drug-susceptible tuberculosis strains. The different colors on the branches indicate different lineages and sublineages. The outside circle indicates treatment outcomes of corresponding patients.

### Identification of a functional mutation set for predicting treatment outcomes

GWAS of the MTB isolates identified fourteen fixed bacterial nonsynonymous variants associated with poor treatment outcomes (Figure 2A). These variants were distributed in thirteen genes involved in intermediary metabolism and respiration (*cobN, dlaT, metA, Rv0648* and *Rv1248c*), cell wall and cell processes (*ctpB, Rv2164c* and *Rv1717*) and virulence (*otsB1* and *Rv3168*), with the *otsB1* G559D mutation showing the strongest association (P=7.3×10^−10^) (Figure 2−table supplement 1).

**Figure 2.**
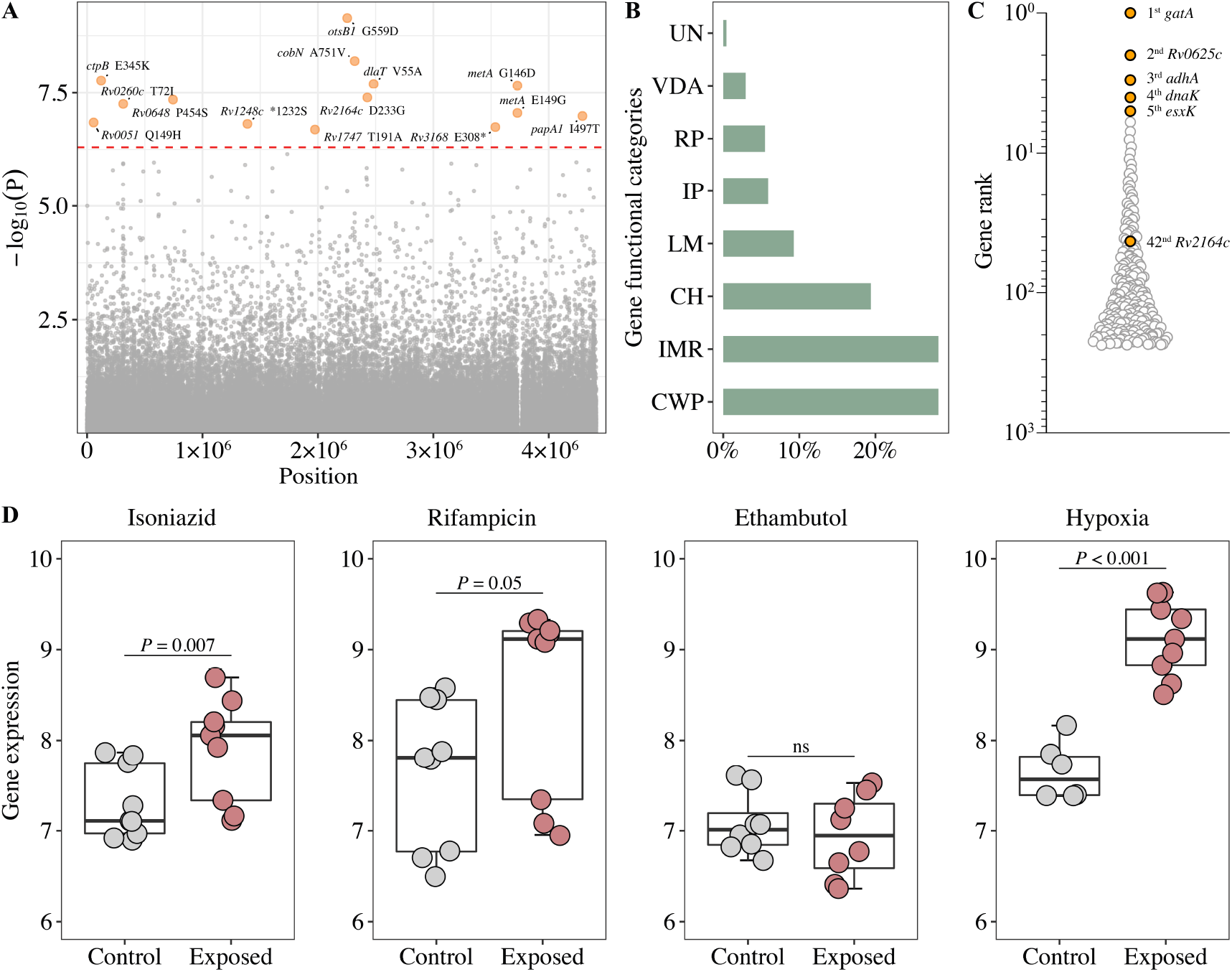
Generation of the functional mutation set. **(A)** Manhattan plots of genome-wide association study (GWAS) for fixed SNPs associated with poor treatment outcomes. The dashed red line highlights the Bonferroni-corrected threshold (P = 5.04×10^−7^). **(B)** Distribution of GWAS identified unfixed SNPs across gene functional categories. CWP: cell wall and cell processes; IMR: intermediary metabolism and respiration; CH: conserved hypotheticals; LM: lipid metabolism; IP: information pathways; RP: regulatory proteins; VDA: virulence, detoxification, adaptation; UN: unknown. **(C)** Gene prioritization strategies (based on P value rank) for significantly associated unfixed SNPs. **(D)** Gene expression from RNA-seq (log_2_FPKM) of *Rv2164c* under drug pressure and hypoxia.

Unfixed mutations are thought to represent adaptive mutations emerging within the host (Nimmo et al., 2020). To investigate whether unfixed mutations affect treatment outcomes, we performed a GWAS analysis of unfixed mutations and found that 237 mutations were associated with poor treatment outcomes (Figure 2−figure supplement 1). The frequency of these mutations concentrated in the range of 5%-10%, and they were predominantly found in genes whose encoded proteins are involved in cell wall and cell processes, intermediary metabolism and respiration (Figure 2B). When the genes carrying unfixed mutations were ranked according to the significance of the mutations’ association with poor outcomes, the highest ranked gene was *gatA*, which has been associated with rifampicin tolerance (Cai et al., 2020) (Figure 2C).

Gene expression patterns under stress conditions can provide important insights into the function of the gene (Bosch et al., 2021). We therefore analyzed the genes containing GWAS-identified fixed mutations for changes in expression after exposure to first-line drugs and hypoxic conditions. This analysis showed that the expression of some of genes increased under these conditions (drug-treated: *Rv2164c, cobN, Rv0260c*; hypoxia: *Rv2164c, otsB1, papA1*), Only one gene, *Rv2164c* (Figure 2D) contained both GWAS-identified fixed (Figure 2A) and unfixed (Figure 2C) mutations, while the expression of others decreased (drug-treated: *otsB1, Rv0648, Rv1248c, Rv3168*; hypoxia: *Rv0648*) (Figure 2D; Figure 2−figure supplement 2), suggesting that perhaps these genes could be involved in adaptation to drug and hypoxic stress.

### Ongoing mutational signatures of ROS associated with TB treatment outcomes

Previous reports have suggested that poor TB treatment outcomes may be associated with increased ROS mutational signatures (C>T/G>A mutations) (Q. Liu et al., 2020; Moreno-Molina et al., 2021). To determine whether the increased ROS signatures were the result of mutations that were fixed before the infection or unfixed because of their *de novo* accumulation during infection, we compared the distribution of six mutation types in fixed and unfixed mutations. In the fixed mutations, there was no significant difference in the proportions of ROS mutational signatures for isolates from patients with good or poor outcomes (43.9% vs. 44.2%, Figure 3A). In the unfixed *de novo* mutations, however, the ROS mutational signatures were significantly more frequent in isolates from patients with poor outcomes (22.9% vs. 26.3%, P=0.027, Figure 3A). A further analysis of the distribution of unfixed mutations across gene functional categories found no difference in the distribution of total unfixed mutations between good and poor outcomes (Figure 3B), but isolates from patients with poor outcomes showed a higher ratio of nonsynonymous mutations (3.9% vs. 5.7%, P=0.048, Figure 3B) in genes belonging to the functional category “information pathway”.

**Figure 3.**
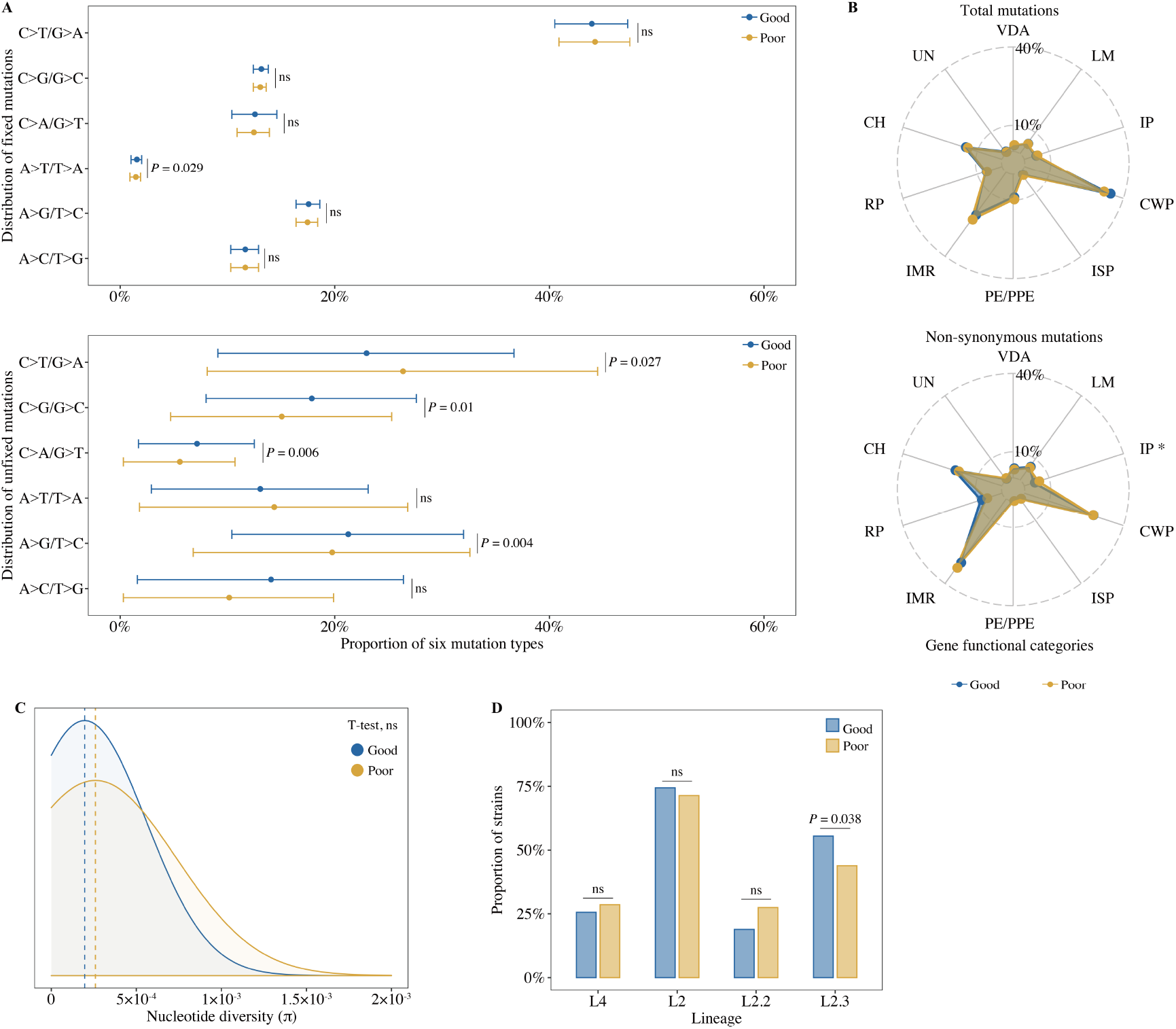
Bacterial whole-genome mutation features between patients with different treatment outcomes. **(A)** The proportion of six mutation types in all fixed and unfixed mutations. **(B)** Distribution of total unfixed mutations and nonsynonymous unfixed mutations across gene functional categories. VDA: virulence, detoxification, adaptation; LM: lipid metabolism; IP: information pathways; CWP: cell wall and cell processes; ISP: insertion seqs and phages; IMR: intermediary metabolism and respiration; RP: regulatory proteins; CH: conserved hypotheticals; UN: unknown. **(C)** Comparison of nucleotide genetic diversity between isolated of patients from with good and poor outcomes. **(D)** Distribution of MTB lineages and sublineages. P-value < 0.05 was considered significant. * P<0.05, ns: no significant.

Nucleotide diversity and the characteristics of the different lineages of MTB are thought to be determinants of virulence and thus may affect TB treatment outcomes. (O’Neill et al., 2015; Tong et al., 2022). An analysis of nucleotide diversity analysis revealed no significant difference in patients with different outcomes (2.0×10^−4^ vs 2.6 ×10^−4^, Figure 3C). In contrast, an analysis of lineage distribution showed that strains belonging to the modern Beijing lineage L2.3, which has been related to increased transmission and high virulence (Tong et al., 2022), were significantly more prevalent in patients with good outcomes (55.5% vs 44.0%, Figure 3D), suggesting that MTB lineage was not associated with poor treatment outcomes in the populations studied.

### GWAS-identified mutations help predict TB treatment outcomes

To identify the risk factors for poor TB treatment outcomes and understand how the variables are related, we used logistic regressions analysis and structural equation models (SEM) with the patients’ clinical characteristics and the bacterial factors associated with poor outcomes. We found that patient age, sex, duration of diagnostic delay, and the GWAS-identified fixed mutations were all associated with poor TB outcomes (Figure 4A). When pooled together, the host factors of age, sex, and duration of diagnostic delay had a larger impact on TB treatment outcomes than GWAS-identified fixed mutations (Figure 4B). Logistic regression, incorporating the identified risk factors, was then used to construct a clinical prediction model that is graphically depicted in a nomogram (Figure 4C). For example, in a 65-year-old male TB patient with a 4-month delay in diagnosis, the risk of poor outcome would increase from 5.4% to 17.4% if his MTB isolate contained at least one GWAS-identified mutation.

**Figure 4.**
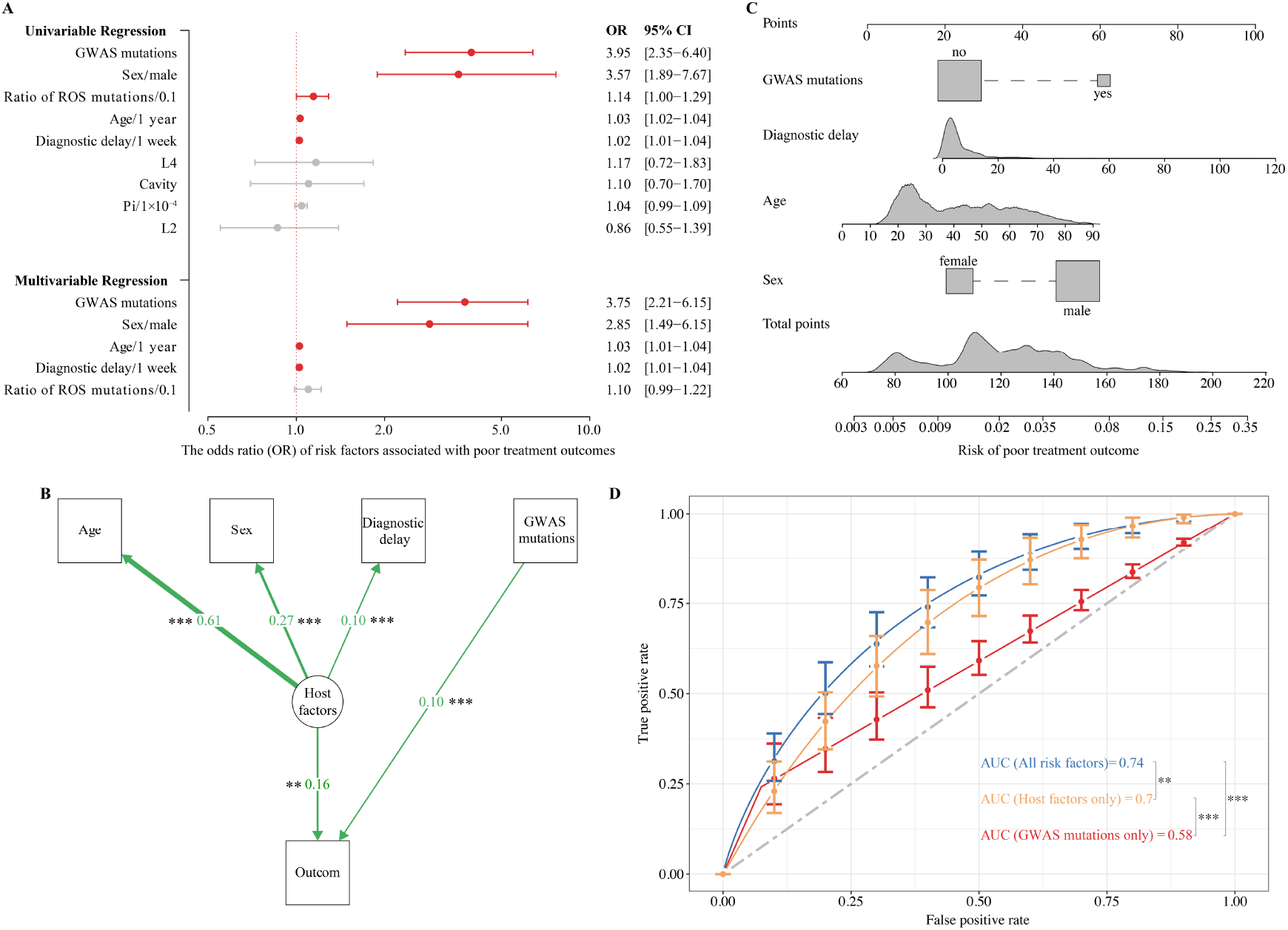
Effects of GWAS identified mutations on tuberculosis treatment outcomes. **(A)** Univariable and multivariable logistic regression on the risk factors for poor treatment outcomes. **(B)** Graphical representation of the structural equation model examining the effect of host and bacterial factors on treatment outcomes. Unidirectional arrows between variables indicate regression and are associated with standardized regression coefficients. **(C)** Nomogram for predicting the probability of poor treatment outcomes. **(D)** ROC curves based on risk factors that may be predictive of tuberculosis treatment outcomes. P-value < 0.05 was considered significant. * P<0.05, **P<0.01, *** P<0.001.

The ROC curve for the clinical prediction model based only on host risk factors had an AUC of 0.70, significantly higher than the AUC of 0.58 (P < 0.001) when only GWAS-identified mutations were included (Figure 4D). However, when both host factors and GWAS-identified mutations were included in the model, the AUC for predicting poor treatment outcomes increased significantly (P = 0.01) from 0.70 to 0.74, with a sensitivity of 0.85 and specificity of 0.55 (Figure 4D). Based on this analysis, it appears that genomic variants of the infecting MTB strain may play a role in determining a patient’s treatment outcome.

## DISCUSSION

To our knowledge, this is the first comprehensive evaluation of the contribution of bacterial genetic factors to poor treatment outcomes in drug-susceptible TB patients. Although the analysis showed that, as expected, patient characteristics of age, sex, and duration of diagnostic delay were associated with poor TB treatment outcomes, it also found fourteen bacterial genomic variants that were also associated with an increased risk of poor treatment outcomes. The best clinical prediction model, with an AUC of 0.74, incorporates both host and bacterial risk factors. In a setting where the genomes of all MTB isolates are sequenced, these risk factors might provide a rationale for the development of personalized treatment approaches.

GWAS has yielded remarkable advances in the understanding of complex traits and has identified hundreds of genetic risk variants in humans (Uffelmann et al., 2021), including genetic polymorphisms associated with increased susceptibility to TB (Quistrebert et al., 2021; Zheng et al., 2018). With the increasing availability of bacterial WGS data has made it possible to use GWAS to study the relationship between pathogen genotypes with clinical disease phenotypes. In MTB, GWAS has identified genetic determinants of drug resistance (Coll et al., 2018; Farhat et al., 2013), TB transmission (Sobkowiak et al., 2020), host preference (Luo et al., 2022) and virulence (Genestet et al., 2022). In this study, GWAS analysis was used to identify bacterial genetic variants associated with poor outcomes in drug-susceptible TB patients, and assess their impact.

However, while this study found that certain mutations in the MTB genome contribute to the risk of poor treatment outcomes, models that only consider bacterial factors are poor predictors of outcome, with an AUC of only 0.58. Furthermore, only 24.2% (22/91) of patients with poor outcomes carried at least one GWAS-identified fixed mutation (Figure 2−table supplement 1), and therefore these fixed mutations we identified played no role in the majority of poor outcomes. The 237 GWAS-identified unfixed mutations are diverse (Figure 2−figure supplement 1), with a mutation frequency of generally of only 5%-10% (Figure 2−figure supplement 3). The low frequency of these diverse unfixed mutations suggests they confer a relatively small selective advantage, without any preferential mutation types or increasing mutation frequency. In addition, none of the GWAS-identified unfixed mutations are among the 14 GWAS-identified fixed mutations, suggesting the unfixed mutations merely reflect individual genetic features of the particular MTB isolate, with no evidence of homoplastic fixation in the bacterial population.

Examining the function of the genes carrying the GWAS-identified mutations could provide insight into their capacity to affect treatment outcomes. Gene *otsB1*, the mutation with the strongest association, encodes trehalose-6-phosphate phosphatase, which has been associated with rifampicin tolerance under hypoxic stress (Jakkala & Ajitkumar, 2019). We also confirmed the previous finding that the expression of *otsB* is up-regulated under hypoxic stress (Jakkala & Ajitkumar, 2019), (Figure 2−figure supplement 2). CtpB is a copper transporting P-type ATPase that helps MTB adapt to non-physiological concentrations of Cu^2+^ inside phagosomes and thereby promotes disease progression by enhancing bacterial survival in macrophages (León-Torres et al., 2020). *metA* encodes homoserine transacetylase, a key enzyme for methionine biosynthesis and considered a potential drug target (Berney et al., 2015; Hasenoehrl et al., 2019). The E2 component of pyruvate dehydrogenase, encoded by *dlaT*, is required for optimal MTB growth and resistance to reactive nitrogen intermediates (RNI) and immune killing by host cells (Shi & Ehrt, 2006; Tian et al., 2005). PapA1 is an acyltransferase essential for the biosynthesis of the MTB virulence factor sulfolipid-1(Kumar et al., 2007). The *cobN* gene encodes a critical protein for the synthesis of vitamin B12, which is essential for MTB proliferation and disease progression (Savvi et al., 2008). Gebremicael et al. (Gebremicael et al., 2019) suggested that the serum concentration of vitamin B12 might be used as a diagnostic biomarker for active TB. Some of the mutations associated with poor outcomes were found in genes of unknown function, such as *Rv2164c, Rv0648* and *Rv3148*, but if their association with treatment outcomes is confirmed in subsequent studies, they might be worth further investigation (Figure 2−figure supplement 2).

While we found a statistically significant effect of the MTB genomic determinants on treatment outcomes of drug sensitive cases, the effect appears to be limited. The 14 GWAS-identified fixed mutations were distributed among only 22 of the 91 patients with poor outcomes and none of the mutations were shared amongst all 22 patients. The mutations may therefore be epistatic adaptations suited to the genomic characteristics of each individual strain. In addition, the GWAS-identified mutations were also present in some patients with good outcomes but were less frequent than in patients with poor outcomes. Although patient information was unavailable concerning other known host risk factors such as adherence to the treatment regimen and comorbidities such as HIV co-infection, diabetes, smoking and low BMI (Kamara et al., 2022; Leung et al., 2015; Vernon et al., 2019), our clinical prediction model still revealed that host factors are significantly more important determinants of poor outcomes than bacterial factors. Nevertheless, we believe that our study has shown that bacterial genomic factors can also contribute to poor outcomes.

Sample size and the classification of treatment outcomes are important challenges when exploring the association of bacterial factors with poor outcomes. Because standard first-line regimens cure at least 85% of drug-susceptible TB patients, this study had to pool data from three sites with a total of 3496 new cases to obtain 91 patients with poor outcomes. We excluded 300 patients who either died from non-TB causes, were lost to follow-up or had unknown outcomes. We attempted to validate our findings with a dataset of 1397 new drug-susceptible TB cases from Malawi (Guerra-Assunção et al., 2015), but we couldn’t replicate the GWAS analysis because the only poor treatment outcome in the Malawi dataset was death, and it was impossible to distinguish the patients who succumbed to TB from those who died from non-TB causes (Figure 2−figure supplement 4).

In conclusion, we found that there are bacterial genomic determinants that significantly increase the risk of poor treatment outcomes in drug-susceptible TB patients. Although host factors are more important, the most accurate models to predict poor treatment outcome with drug-susceptible TB incorporate both host and bacterial risk factors. In the future, it may be possible to identify patients at high risk for treatment failure by analyzing the characteristics of the host and the bacterial genome of the infecting strain. This could make it possible to identify patients requiring longer or individualized treatment regimens and thus improve cure rates for drug-susceptible TB.

## MATERIALS AND METHODS

### Selection of patients and samples

A strain database search was performed for TB patients treated during 2009-2020 at three study sites in Shanghai, Sichuan, and Heilongjiang, China. For each of the 4374 TB patients registered during this period, pretreatment isolates for each patient were collected for WGS, WGS data and the patients’ demographic and clinical features were obtained from a published study (Li et al., 2022). All new cases susceptible to first-line drugs (rifampicin, isoniazid, pyrazinamide, ethambutol) by genotypic drug-susceptibility testing (gDST), and whose records contained treatment outcomes, were selected for the study.

The WHO recommended treatment outcome definitions for TB are cured, treatment completed, treatment failed, died, lost to follow-up and not evaluated (Linh et al., 2021). Of these, patients who died were divided into deaths from TB and non-TB, and those not evaluated included cases transferred for treatment of multidrug-resistant tuberculosis (MDR-TB) and cases whose treatment outcome was unknown. For the current study, TB treatment outcomes were grouped into 3 categories: 1) good outcomes -- cured and treatment completed; 2) poor outcomes -- treatment failures, deaths from TB, transferred for MDR and relapse; 3) other -- lost to follow-up, non-TB deaths and unknown outcome.

### SNPs calling, resistance prediction and phylogenetic reconstruction

A previously described pipeline was used for calling single nucleotide polymorphisms (SNPs) (Chen et al., 2021). Briefly, the Sickle tool was used to trim WGS data to retain reads with a Phred base quality above 20 and length greater than 30 nucleotides. Reads were mapped to the MTB H37Rv reference strain (GenBank AL123456) with bowtie2 (v2.2.9), and then SAMtools (v1.3.1) was used for SNP-calling with a mapping quality greater than 30. Varscan (v2.3.9) was used to identified fixed (frequency, ≥ 75%) and unfixed (< 75%) SNPs with at least 5 supporting reads and the strand bias filter option on. The drug-resistance profile and lineages were predicted from WGS data using SAM-TB (Yang et al., 2022). Phylogenetic trees were constructed using the maximum-likelihood method (RAxML-NG) (Kozlov et al., 2019) and visualized on the Interactive Tree of Life platform (https://itol.embl.de/).

### Estimates of nucleotide diversity and GWAS analyses

Nucleotide diversity (π) was estimated using the PoPoolation package (Kofler et al., 2011). Following O’Neill et al. (O’Neill et al., 2015), we randomly subsampled (n = 10) read data from each sample to a uniform 50× coverage to limit the effects of differential coverage across samples. Using the subsampled data with uniform coverage, we then calculated nucleotide diversity in 100 kb sliding windows across the genome in 10 kb steps. GWAS analyses were performed using GEMMA software (v0.98.3) (Zhou & Stephens, 2012) to identify nonsynonymous variants associated with poor TB treatment outcomes. A linear mixed model was used to control for the confounding effects of MTB lineage, sublineage and outbreak-based population structure (Coll et al., 2018), and the significance threshold was adjusted with the Bonferroni correction.

### RNA-seq data collection and analysis

Raw RNA-Seq read data (GSE165581: INH, GSE166622: RIF, GSE118084: EMB, and GSE116353: hypoxia) from MTB laboratory strain H37Rv exposed to first-line drugs and hypoxic conditions were downloaded from the Gene Expression Omnibus (GEO) database (https://www.ncbi.nlm.nih.gov/geo/). Sequencing reads passing quality control were aligned to the MTB H37Rv reference strain using bowtie2. Unique reads were selected and sorted using SAMtools, then quantitated using htseq-count (v0.11.3). FPKM values calculated by DESeq2 (v1.26.0) were used as measures of gene expression, and genes with |log_2_(fold change)| ≥1 and p values < 0.05 were considered differentially expressed.

### Statistical analysis

The t-test was used for comparing nucleotide diversity and the ratios of the six mutation types between TB patients with good and poor treatment outcomes. The chisquare test was used to assess whether the distribution of MTB lineages and mutations across gene functional categories in TubercuList (Lew et al., 2011) differed between patients with different treatment outcomes. Factors associated with poor treatment outcomes were tested with logistic regression in univariate and multivariate analyses. Structural equation models (SEM) were generated to determine the structural relation between the observed variables and latent constructs (Giam & Olden, 2016). We constructed logistic regression models with the selected bacterial and host factors as predictors of TB treatment outcome, and the ROC curves of the prediction models were compared using DeLong’s test.

## DECLARATIONS

## Acknowledgements

We thank Dr. Mia Crampin, Dr. Judith Glynn and Ms Estelle McLean from London School of Hygiene and Tropical Medicine for supplying or collating patient information of Malawi dataset, and Dr. Qingyun Liu from Harvard T. H. Chan School of Public Health for the constructive comments provided on an earlier version of this paper.

## Authors’ contributions

All authors designed the paper by discussing the key concerns to be include. YC and QG designed research. YC, QJ, MP and QG performed research. YC, HET and QG wrote the paper. All authors approved the final draft.

## Data availability

Files containing sequencing reads were deposited in the National Institutes of Health Sequence Read Archive under BioProject PRJNA869190.

## Supplementary Materials

**Figure 2-table supplement 1.**
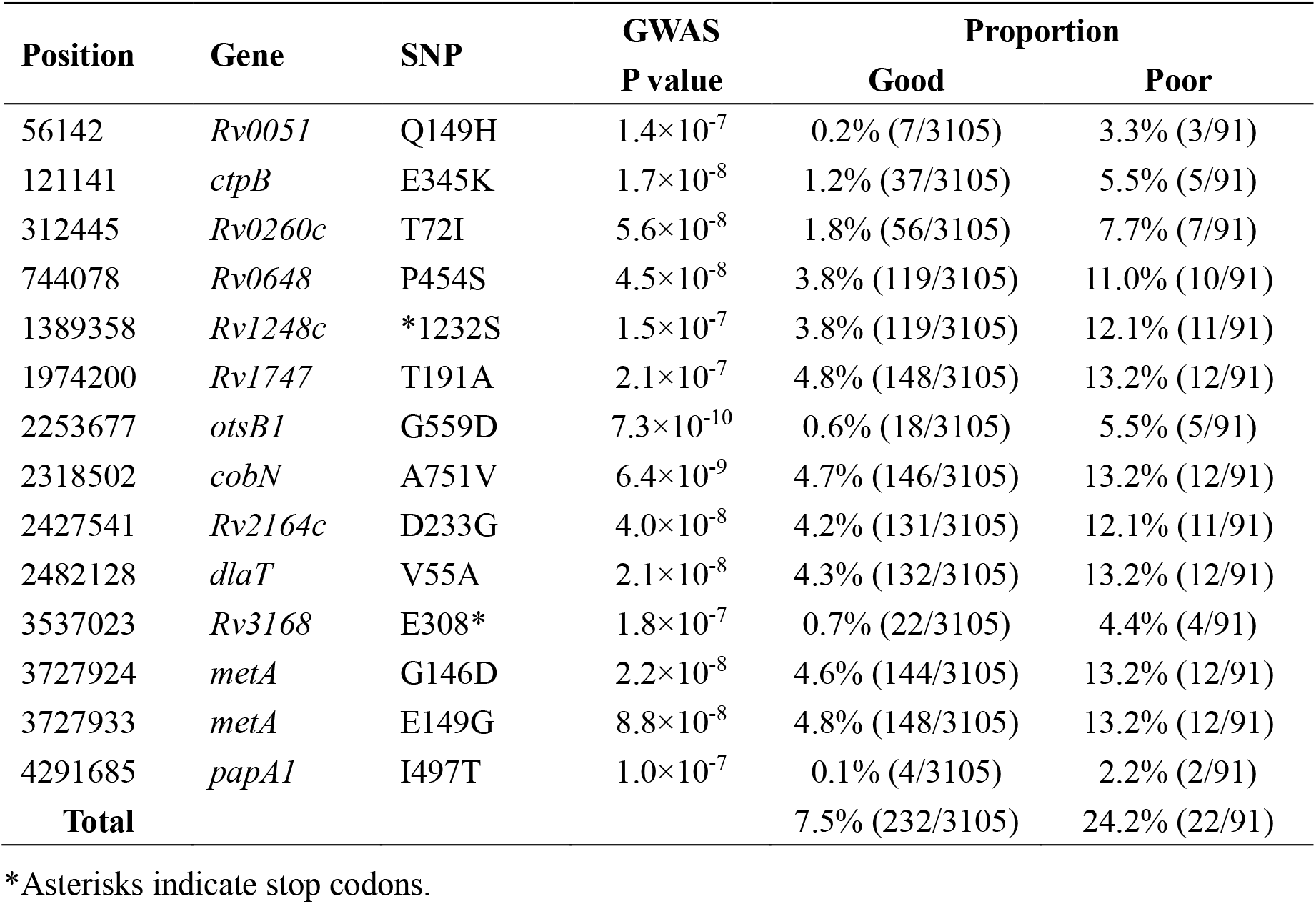
GWAS identified fixed SNPs.

**Figure 2-figure supplement 1.**
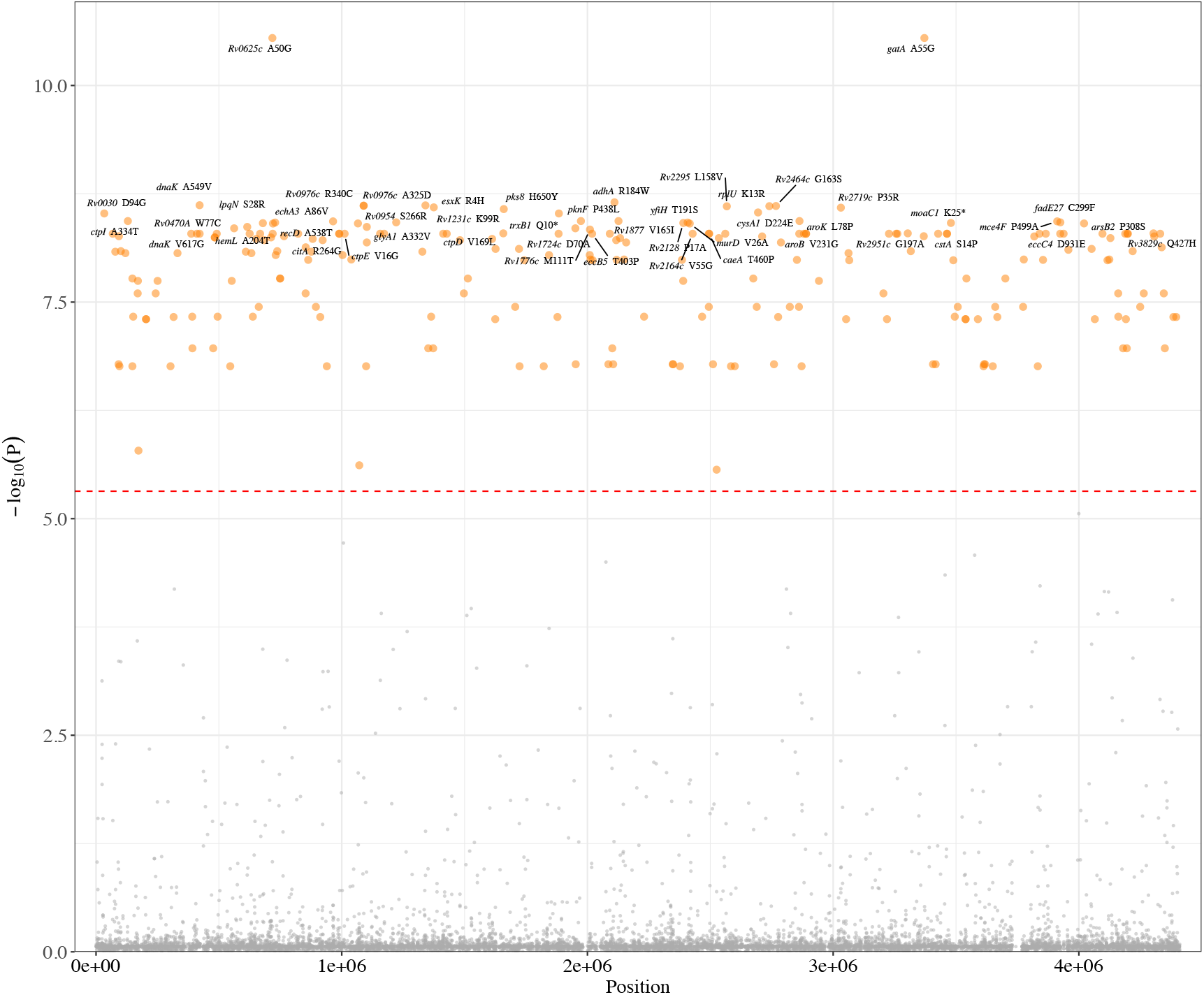
Manhattan plots of unfixed SNPs associated with poor treatment outcomes. The top 50 unfixed mutations were annotated with the gene. The dashed red line highlights the Bonferroni-corrected threshold (P = 4.82×10^−6^)

**Figure 2-figure supplement 2.**
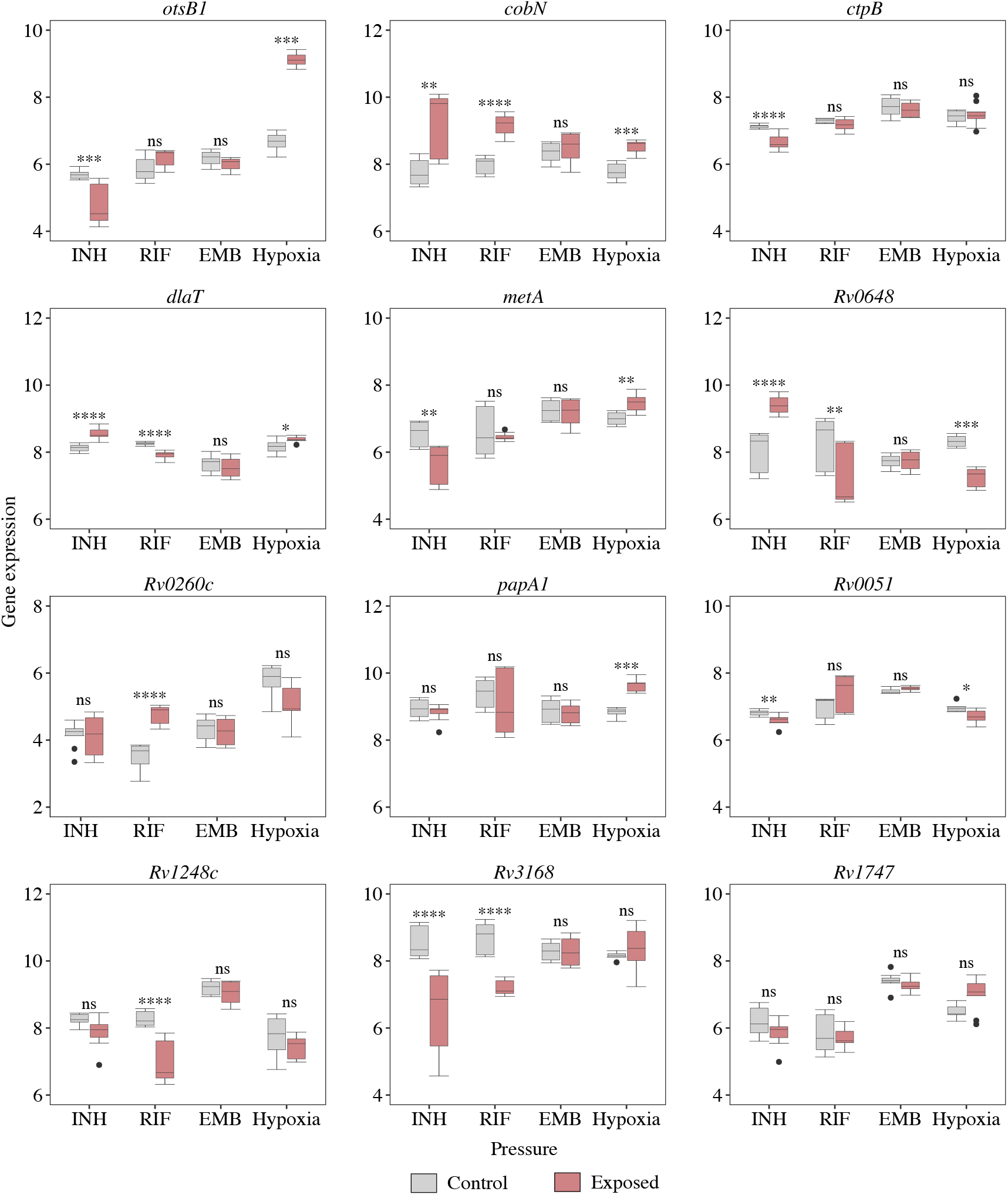
Gene expression (log_2_FPKM) from RNA-seq after drug exposure and hypoxia.

**Figure 2-figure supplement 3.**
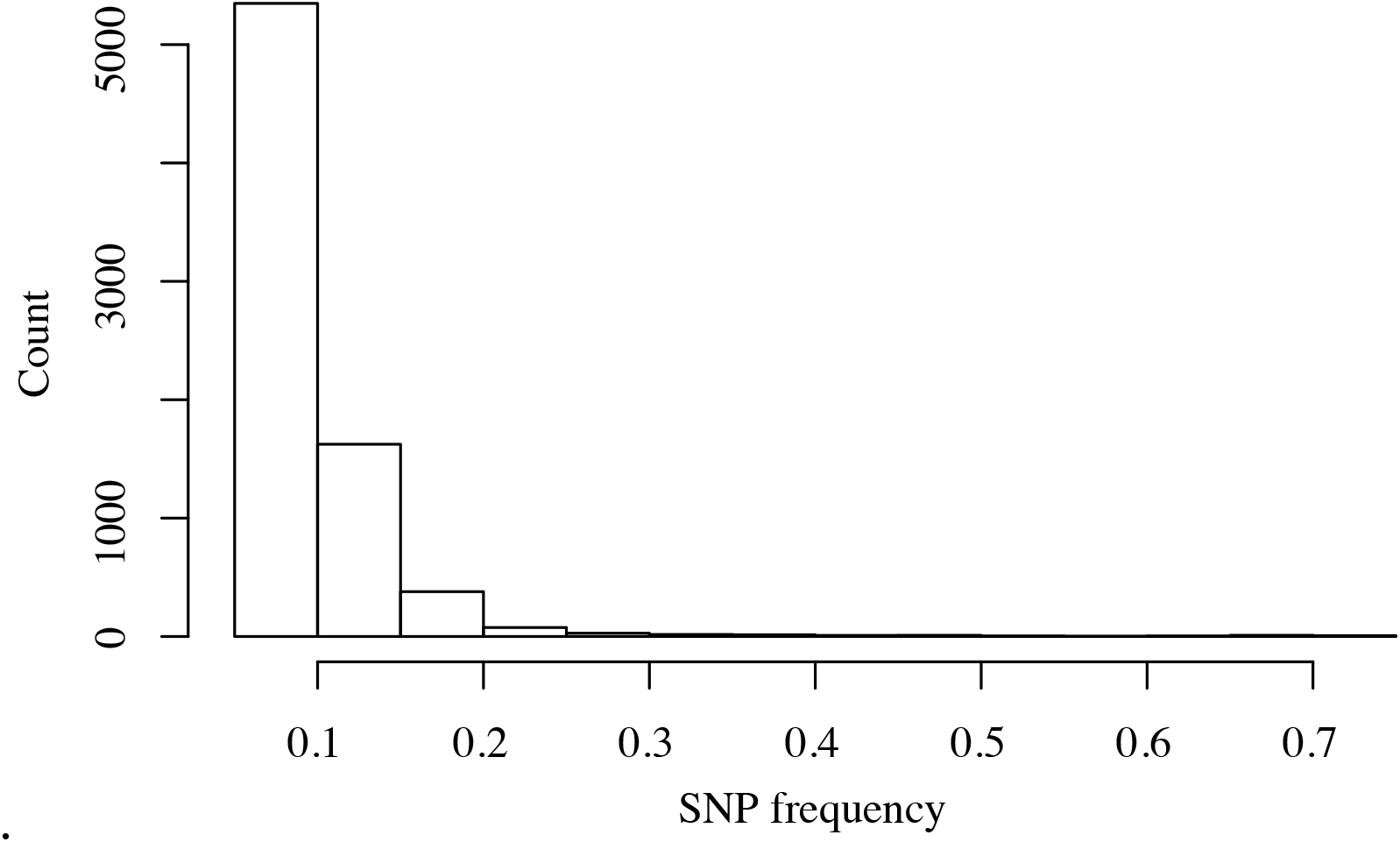
Within-host frequency distribution of GWAS-identified unfixed mutations.

**Figure 2-figure supplement 4.**
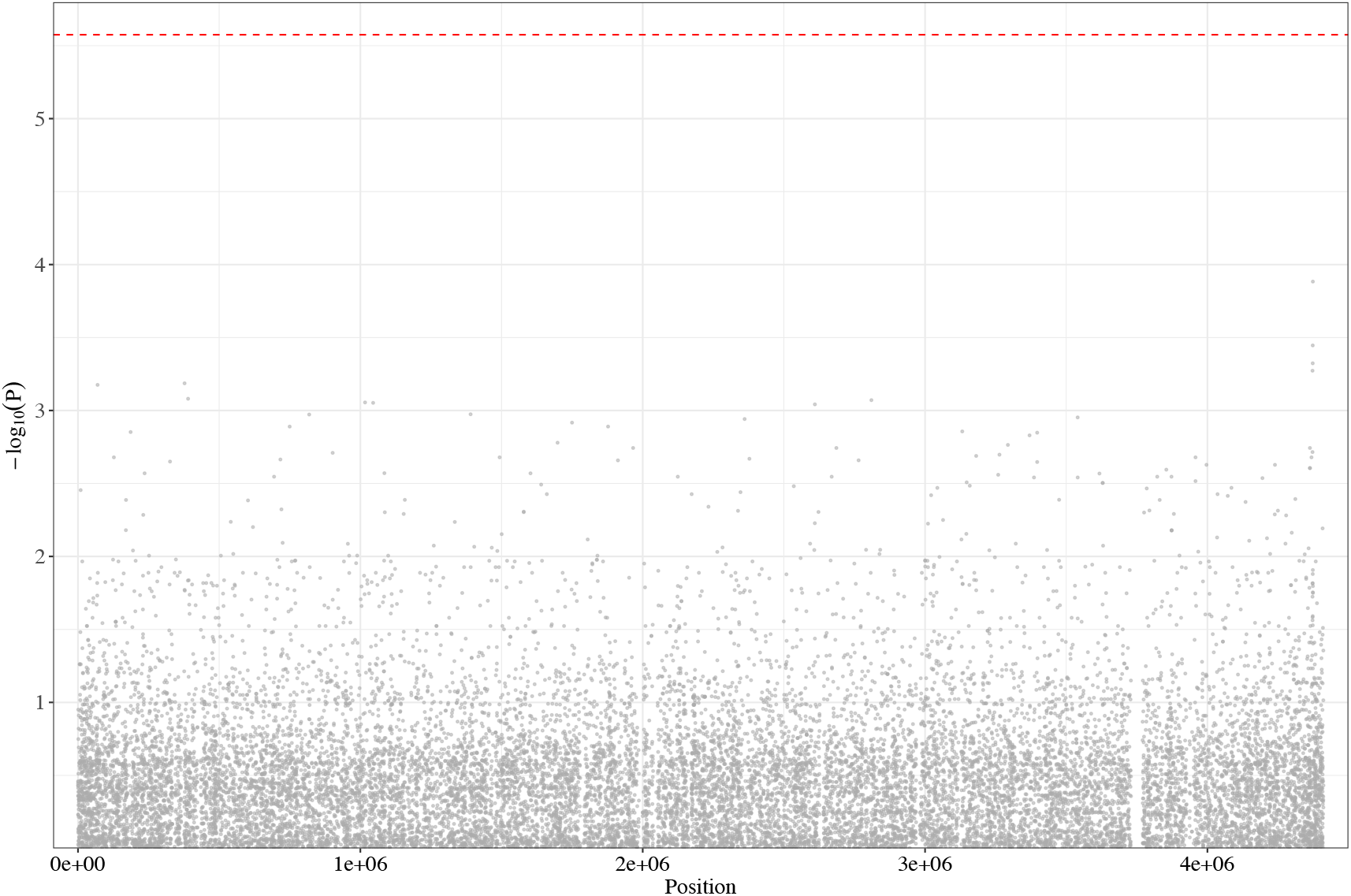
Manhattan plot of GWAS analysis based on the Malawi dataset. The dashed red line highlights the Bonferroni-corrected threshold (P = 2.66×10^−6^)

## Reference

Alipanah N, Jarlsberg L, Miller C, Linh NN, Falzon D, Jaramillo E, Nahid P (2018) Adherence interventions and outcomes of tuberculosis treatment: A systematic review and meta-analysis of trials and observational studies Plos Medicine 15:e1002595. https://doi.org/10.1371/journal.pmed.1002595

Bastos ML, Lan Z, Menzies D (2017) An updated systematic review and meta-analysis for treatment of multidrug-resistant tuberculosis European Respiratory Journal 49:1600803. https://doi.org/10.1183/13993003.00803-2016

Berney M, Berney-Meyer L, Wong KW, Chen B, Chen M, Kim J, Wang J, Harris D, Parkhill J, Chan J, Wang F, Jacobs WR Jr (2015) Essential roles of methionine and S-adenosylmethionine in the autarkic lifestyle of Mycobacterium tuberculosis Proceedings of the National Academy of Sciences 112:10008–10013. https://doi.org/10.1073/pnas.1513033112

Bosch B, DeJesus MA, Poulton NC, Zhang W, Engelhart CA, Zaveri A, Lavalette S, Ruecker N, Trujillo C, Wallach JB, Li S, Ehrt S, Chait BT, Schnappinger D, Rock JM (2021) Genome-wide gene expression tuning reveals diverse vulnerabilities of M. tuberculosis Cell 184:4579–4592 e4524. https://doi.org/10.1016/j.cell.2021.06.033

Brauner A, Fridman O, Gefen O, Balaban NQ (2016) Distinguishing between resistance, tolerance and persistence to antibiotic treatment Nature Reviews Microbiology 14:320–330. https://doi.org/10.1038/nrmicro.2016.34

Cai RJ, Su HW, Li YY, Javid B (2020) Forward Genetics Reveals a gatC-gatA Fusion Polypeptide Causes Mistranslation and Rifampicin Tolerance in Mycobacterium smegmatis Frontiers in Microbiology 11:577756. https://doi.org/10.3389/fmicb.2020.577756

Chen YW, Ji LC, Liu QY, Li JL, Hong CY, Jiang Q, Gan MY, Takiff HE, Yu WY, Tan WG, Gao Q (2021) Lesion Heterogeneity and Long-Term Heteroresistance in Multidrug-Resistant Tuberculosis The Journal of Infectious Diseases 224:889–893. https://doi.org/10.1093/infdis/jiab011

Chenciner L, Annerstedt KS, Pescarini JM, Wingfield T (2021) Social and health factors associated with unfavourable treatment outcome in adolescents and young adults with tuberculosis in Brazil: a national retrospective cohort study The Lancet Global Health 9:e1380–e1390. https://doi.org/10.1016/S2214-109X(21)00300-4

Coll F, Phelan J, Hill-Cawthorne GA, Nair MB, Mallard K, Ali S, Abdallah AM, Alghamdi S, Alsomali M, Ahmed AO, Portelli S, Oppong Y, Alves A, Bessa TB, Campino S, Caws M, Chatterjee A, Crampin AC, Dheda K, Furnham N, Glynn JR, Grandjean L, Minh Ha D, Hasan R, Hasan Z, Hibberd ML, Joloba M, Jones-Lopez EC, Matsumoto T, Miranda A, Moore DJ, Mocillo N, Panaiotov S, Parkhill J, Penha C, Perdigao J, Portugal I, Rchiad Z, Robledo J, Sheen P, Shesha NT, Sirgel FA, Sola C, Oliveira Sousa E, Streicher EM, Helden PV, Viveiros M, Warren RM, McNerney R, Pain A, Clark TG (2018) Genome-wide analysis of multi- and extensively drug-resistant Mycobacterium tuberculosis Nature Genetics 50:307–316. https://doi.org/10.1038/s41588-017-0029-0

Dheda K, Gumbo T, Maartens G, Dooley KE, McNerney R, Murray M, Furin J, Nardell EA, London L, Lessem E, Theron G, van Helden P, Niemann S, Merker M, Dowdy D, Van Rie A, Siu GK, Pasipanodya JG, Rodrigues C, Clark TG, Sirgel FA, Esmail A, Lin HH, Atre SR, Schaaf HS, Chang KC, Lange C, Nahid P, Udwadia ZF, Horsburgh CR, Jr., Churchyard GJ, Menzies D, Hesseling AC, Nuermberger E, McIlleron H, Fennelly KP, Goemaere E, Jaramillo E, Low M, Jara CM, Padayatchi N, Warren RM (2017) The epidemiology, pathogenesis, transmission, diagnosis, and management of multidrug-resistant, extensively drug-resistant, and incurable tuberculosis The lancet Respiratory medicine 5:291–360. https://doi.org/10.1016/S2213-2600(17)30079-6

Farhat MR, Shapiro BJ, Kieser KJ, Sultana R, Jacobson KR, Victor TC, Warren RM, Streicher EM, Calver A, Sloutsky A, Kaur D, Posey JE, Plikaytis B, Oggioni MR, Gardy JL, Johnston JC, Rodrigues M, Tang PK, Kato-Maeda M, Borowsky ML, Muddukrishna B, Kreiswirth BN, Kurepina N, Galagan J, Gagneux S, Birren B, Rubin EJ, Lander ES, Sabeti PC, Murray M (2013) Genomic analysis identifies targets of convergent positive selection in drug-resistant Mycobacterium tuberculosis Nature Genetics 45:1183–1189. https://doi.org/10.1038/ng.2747

Gebremicael G, Alemayehu M, Sileshi M, Geto Z, Gebreegziabxier A, Tefera H, Ashenafi N, Tadese C, Wolde M, Kassa D (2019) The serum concentration of vitamin B12 as a biomarker of therapeutic response in tuberculosis patients with and without human immunodeficiency virus (HIV) infection International Journal of General Medicine 12:353–361. https://doi.org/10.2147/IJGM.S218799

Genestet C, Refrégier G, Hodille E, Zein-Eddine R, Le Meur A, Hak F, Barbry A, Westeel E, Berland J-L, Engelmann A, Verdier I, Lina G, Ader F, Dray S, Jacob L, Massol F, Venner S, Dumitrescu O (2022) Mycobacterium tuberculosis genetic features associated with pulmonary tuberculosis severity International Journal of Infectious Diseases 125:74–83. https://doi.org/10.1016/j.ijid.2022.10.026

Giam X, Olden JD (2016) Quantifying variable importance in a multimodel inference framework Methods in Ecology and Evolution 7:388–397. https://doi.org/10.1111/2041-210X.12492

Guerra-Assunção J, Crampin A, Houben R, Mzembe T, Mallard K, Coll F, Khan P, Banda L, Chiwaya A, Pereira R (2015) Large-scale whole genome sequencing of M. tuberculosis provides insights into transmission in a high prevalence area Elife 4:e05166. https://doi.org/10.7554/eLife.05166

Hasenoehrl EJ, Rae Sajorda D, Berney-Meyer L, Johnson S, Tufariello JM, Fuhrer T, Cook GM, Jacobs WR, Jr., Berney M (2019) Derailing the aspartate pathway of Mycobacterium tuberculosis to eradicate persistent infection Nature Communications 10:4215. https://doi.org/10.1038/s41467-019-12224-3

Hicks ND, Yang J, Zhang X, Zhao B, Grad YH, Liu L, Ou X, Chang Z, Xia H, Zhou Y, Wang S, Dong J, Sun L, Zhu Y, Zhao Y, Jin Q, Fortune SM (2018) Clinically prevalent mutations in Mycobacterium tuberculosis alter propionate metabolism and mediate multidrug tolerance Nature Microbiology 3:1032–1042. https://doi.org/10.1038/s41564-018-0218-3

Imperial MZ, Nahid P, Phillips PPJ, Davies GR, Fielding K, Hanna D, Hermann D, Wallis RS, Johnson JL, Lienhardt C, Savic RM (2018) A patient-level pooled analysis of treatment-shortening regimens for drug-susceptible pulmonary tuberculosis Nature Medicine 24:1708–1715. https://doi.org/10.1038/s41591-018-0224-2

Jakkala K, Ajitkumar P (2019) Hypoxic non-replicating persistent Mycobacterium tuberculosis develops thickened outer layer that helps in restricting rifampicin entry Frontiers in Microbiology 10:2339. https://doi.org/10.3389/fmicb.2019.02339

Kamara RF, Saunders MJ, Sahr F, Losa-Garcia JE, Foray L, Davies G, Wingfield T (2022) Social and health factors associated with adverse treatment outcomes among people with multidrug-resistant tuberculosis in Sierra Leone: a national, retrospective cohort study The Lancet Global Health 10:e543–e554. https://doi.org/10.1016/S2214-109X(22)00004-3

Kofler R, Orozco-terWengel P, De Maio N, Pandey RV, Nolte V, Futschik A, Kosiol C, Schlotterer C (2011) PoPoolation: a toolbox for population genetic analysis of next generation sequencing data from pooled individuals Plos One 6:e15925. https://doi.org/10.1371/journal.pone.0015925

Kozlov AM, Darriba D, Flouri T, Morel B, Stamatakis A (2019) RAxML-NG: a fast, scalable and user-friendly tool for maximum likelihood phylogenetic inference Bioinformatics 35:4453–4455. https://doi.org/10.1093/bioinformatics/btz305

Kumar P, Schelle MW, Jain M, Lin FL, Petzold CJ, Leavell MD, Leary JA, Cox JS, Bertozzi CR (2007) PapA1 and PapA2 are acyltransferases essential for the biosynthesis of the Mycobacterium tuberculosis virulence factor sulfolipid-1 Proceedings of the National Academy of Sciences 104:11221–11226. https://doi.org/10.1073/pnas.0611649104

Lange C, Dheda K, Chesov D, Mandalakas AM, Udwadia Z, Horsburgh Jr CR (2019) Management of drug-resistant tuberculosis The Lancet 394:953–966. https://doi.org/10.1016/S0140-6736(19)31882-3

León-Torres A, Arango E, Castillo E, Soto CY (2020) CtpB is a plasma membrane copper (I) transporting P-type ATPase of Mycobacterium tuberculosis Biological research 53:1–13. https://doi.org/10.1186/s40659-020-00274-7

Lestari BW, McAllister S, Hadisoemarto PF, Afifah N, Jani ID, Murray M, van Crevel R, Hill PC, Alisjahbana B (2020) Patient pathways and delays to diagnosis and treatment of tuberculosis in an urban setting in Indonesia The Lancet Regional Health-Western Pacific 5:100059. https://doi.org/10.1016/j.lanwpc.2020.100059

Leung CC, Yew WW, Chan CK, Chang KC, Law WS, Lee SN, Tai LB, Leung EC, Au RK, Huang SS, Tam CM (2015) Smoking adversely affects treatment response, outcome and relapse in tuberculosis European Respiratory Journal 45:738–745. https://doi.org/10.1183/09031936.00114214

Lew JM, Kapopoulou A, Jones LM, Cole ST (2011) TubercuList--10 years after Tuberculosis (Edinb) 91:1–7. https://doi.org/10.1016/j.tube.2010.09.008

Li M, Guo M, Peng Y, Jiang Q, Xia L, Zhong S, Qiu Y, Su X, Zhang S, Yang C, Mijiti P, Mao Q, Takiff H, Li F, Chen C, Gao Q (2022) High proportion of tuberculosis transmission among social contacts in rural China: a 12-year prospective population-based genomic epidemiological study Emerging Microbes & Infections 11:2102–2111. https://doi.org/10.1080/22221751.2022.2112912

Linh NN, Viney K, Gegia M, Falzon D, Glaziou P, Floyd K, Timimi H, Ismail N, Zignol M, Kasaeva T, Mirzayev F (2021) World Health Organization treatment outcome definitions for tuberculosis: 2021 update European Respiratory Journal 58:2100804. https://doi.org/10.1183/13993003.00804-2021

Liu JF, Gefen O, Ronin I, Bar-Meir M, Balaban NQ (2020) Effect of tolerance on the evolution of antibiotic resistance under drug combinations Science 367:200–204. https://doi.org/10.1126/science.aay3041

Liu Q, Wei J, Li Y, Wang M, Su J, Lu Y, Lopez MG, Qian X, Zhu Z, Wang H, Gan M, Jiang Q, Fu YX, Takiff HE, Comas I, Li F, Lu X, Fortune SM, Gao Q (2020) Mycobacterium tuberculosis clinical isolates carry mutational signatures of host immune environments Science Advances 6:eaba4901. https://doi.org/10.1126/sciadv.aba4901

Liu Q, Zhu J, Dulberger CL, Stanley S, Wilson S, Chung ES, Wang X, Culviner P, Liu YJ, Hicks ND (2022) Tuberculosis treatment failure associated with evolution of antibiotic resilience bioRxiv. https://doi.org/10.1101/2022.03.29.486233

Luo Y, Huang C-C, Liu Q, Howard N, Li X, Zhu J, Amariuta T, Asgari S, Ishigaki K, Calderon R (2022) A FLOT1 host regulatory allele is associated with a recently expanded Mtb clade in patients with tuberculosis medRxiv. https://doi.org/10.1101/2022.02.07.22270622

Migliori GB, Nardell E, Yedilbayev A, D’Ambrosio L, Centis R, Tadolini M, Van Den Boom M, Ehsani S, Sotgiu G, Dara M (2019) Reducing tuberculosis transmission: a consensus document from the World Health Organization Regional Office for Europe European Respiratory Journal 53:1900391. https://doi.org/10.1183/13993003.00391-2019

Mirzayev F, Viney K, Linh NN, Gonzalez-Angulo L, Gegia M, Jaramillo E, Zignol M, Kasaeva T (2021) World Health Organization recommendations on the treatment of drug-resistant tuberculosis, 2020 update European Respiratory Journal 57. https://doi.org/10.1183/13993003.03300-2020

Moreno-Molina M, Shubladze N, Khurtsilava I, Avaliani Z, Bablishvili N, Torres-Puente M, Villamayor L, Gabrielian A, Rosenthal A, Vilaplana C, Gagneux S, Kempker RR, Vashakidze S, Comas I (2021) Genomic analyses of Mycobacterium tuberculosis from human lung resections reveal a high frequency of polyclonal infections Nature Communications 12:2716. https://doi.org/10.1038/s41467-021-22705-z

Nimmo C, Brien K, Millard J, Grant AD, Padayatchi N, Pym AS, O’Donnell M, Goldstein R, Breuer J, Balloux F (2020) Dynamics of within-host Mycobacterium tuberculosis diversity and heteroresistance during treatment Ebiomedicine 55:102747. https://doi.org/10.1016/j.ebiom.2020.102747

O’Neill MB, Mortimer TD, Pepperell CS (2015) Diversity of Mycobacterium tuberculosis across Evolutionary Scales Plos Pathogens 11:e1005257. https://doi.org/10.1371/journal.ppat.1005257

World Health Organization. (2021). Global tuberculosis report 2021. World Health Organization.

Quistrebert J, Orlova M, Kerner G, Ton LT, Luong NT, Danh NT, Vincent QB, Jabot-Hanin F, Seeleuthner Y, Bustamante J (2021) Genome-wide association study of resistance to Mycobacterium tuberculosis infection identifies a locus at 10q26. 2 in three distinct populations Plos Genetics 17:e1009392. https://doi.org/10.1371/journal.pgen.1009392

Savvi S, Warner DF, Kana BD, McKinney JD, Mizrahi V, Dawes SS (2008) Functional characterization of a vitamin B12-dependent methylmalonyl pathway in Mycobacterium tuberculosis: implications for propionate metabolism during growth on fatty acids Journal of Bacteriology 190:3886–3895. https://doi.org/10.1128/JB.01767-07

Shi S, Ehrt S (2006) Dihydrolipoamide acyltransferase is critical for Mycobacterium tuberculosis pathogenesis Infection and immunity 74:56–63. https://doi.org/10.1128/IAI.74.1.56-63.2006

Sobkowiak B, Banda L, Mzembe T, Crampin AC, Glynn JR, Clark TG (2020) Bayesian reconstruction of Mycobacterium tuberculosis transmission networks in a high incidence area over two decades in Malawi reveals associated risk factors and genomic variants Microbial Genomics 6:e000361. https://doi.org/10.1099/mgen.0.000361

Tian J, Bryk R, Shi S, Erdjument-Bromage H, Tempst P, Nathan C (2005) Mycobacterium tuberculosis appears to lack alpha-ketoglutarate dehydrogenase and encodes pyruvate dehydrogenase in widely separated genes Molecular Microbiology 57:859–868. https://doi.org/10.1111/j.1365-2958.2005.04741.x

Tong J, Meng L, Bei C, Liu Q, Wang M, Yang T, Takiff HE, Zhang S, Gao Q, Wang C (2022) Modern Beijing sublineage of Mycobacterium tuberculosis shift macrophage into a hyperinflammatory status Emerging Microbes & Infections 11:715–724. https://doi.org/10.1080/22221751.2022.2037395

Torrey HL, Keren I, Via LE, Lee JS, Lewis K (2016) High Persister Mutants in Mycobacterium tuberculosis Plos One 11:e0155127. https://doi.org/10.1371/journal.pone.0155127

Uffelmann E, Huang QQ, Munung NS, De Vries J, Okada Y, Martin AR, Martin HC, Lappalainen T, Posthuma D (2021) Genome-wide association studies Nature Reviews Methods Primers 1:1–21. https://doi.org/10.1038/s43586-021-00056-9

Vernon A, Fielding K, Savic R, Dodd L, Nahid P (2019) The importance of adherence in tuberculosis treatment clinical trials and its relevance in explanatory and pragmatic trials Plos Medicine 16:e1002884. https://doi.org/10.1371/journal.pmed.1002884

Yang T, Gan M, Liu Q, Liang W, Tang Q, Luo G, Zuo T, Guo Y, Hong C, Li Q (2022) SAM-TB: a whole genome sequencing data analysis website for detection of Mycobacterium tuberculosis drug resistance and transmission Briefings in bioinformatics 23:bbac030. https://doi.org/10.1093/bib/bbac030

Zheng R, Li Z, He F, Liu H, Chen J, Chen J, Xie X, Zhou J, Chen H, Wu X, Wu J, Chen B, Liu Y, Cui H, Fan L, Sha W, Liu Y, Wang J, Huang X, Zhang L, Xu F, Wang J, Feng Y, Qin L, Yang H, Liu Z, Cui Z, Liu F, Chen X, Gao S, Sun S, Shi Y, Ge B (2018) Genome-wide association study identifies two risk loci for tuberculosis in Han Chinese Nature Communications 9:4072. https://doi.org/10.1038/s41467-018-06539-w

Zhou X, Stephens M (2012) Genome-wide efficient mixed-model analysis for association studies Nature Genetics 44:821–824. https://doi.org/10.1038/ng.2310

